# Effects of isoflurane and urethane anesthetics on glutamate neurotransmission in rat brain using *in vivo* amperometry

**DOI:** 10.1101/2023.02.16.528856

**Authors:** Joshua A. Beitchman, Gokul Krishna, Caitlin E. Bromberg, Theresa Currier Thomas

## Abstract

Aspects of glutamate neurotransmission implicated in normal and pathological conditions are often evaluated using *in vivo* recording paradigms in rats anesthetized with isoflurane or urethane. Urethane and isoflurane anesthesia influence glutamate neurotransmission through different mechanisms; however real-time outcome measures of potassium chloride (KCl)-evoked glutamate overflow and glutamate clearance kinetics have not been compared within and between regions of the brain. In the following experiments, *in vivo* amperometric recordings of KCl-evoked glutamate overflow and glutamate clearance kinetics (uptake rate and T_80_) in the cortex, hippocampus and thalamus were performed using glutamate-selective microelectrode arrays (MEAs) in young adult male, Sprague-Dawley rats anesthetized with isoflurane or urethane. Potassium chloride (KCl)-evoked glutamate overflow was similar under urethane and isoflurane anesthesia in all brain regions studied. Analysis of glutamate clearance determined that the uptake rate was significantly faster (53.2%, p<0.05) within the thalamus under urethane compared to isoflurane, but no differences were measured in the cortex or hippocampus. Under urethane, glutamate clearance parameters were region dependent, with significantly faster glutamate clearance in the thalamus compared to the cortex but not the hippocampus (p<0.05). No region dependent differences were measured for glutamate overflow using isoflurane. These data support that amperometric recordings of glutamate under isoflurane and urethane anesthesia result in mostly similar and comparable data. However, certain parameters of glutamate uptake vary based on choice of anesthesia and brain region. Special considerations must be given to these areas when considering comparison to previous literature and when planning future experiments.

**Highlights:**

- KCl-evoked glutamate overflow was similar under urethane and isoflurane anesthesia
- Glutamate clearance parameters were similar under both anesthesias in the cortex and hippocampus but not the thalamus
- Glutamate clearance kinetics differ between brain regions when anesthetized with urethane.
- Experimental design, brain region of interest, and outcome parameters of glutamate clearance should be considered when designing anesthetized amperometry recordings of glutamate.

## INTRODUCTION

Amperometric techniques have provided fundamental information regarding brain chemical communication of neurotransmitter systems in normal and disease-related physiologies. Glutamate is the primary excitatory neurotransmitter in the central nervous system (CNS) [1]. Experimental models of many CNS disorders, including Alzheimer’s disease (AD), Huntington’s disease (HD), Parkinson’s disease (PD), epilepsy, attention-deficit/hyperactivity disorder (ADHD), post-concussive symptoms, and schizophrenia have indicated changes in glutamate neurotransmission as part of their pathophysiology. Mechanisms controlling glutamate neurotransmission provide novel targets for pharmacological intervention to restore physiological norms.

Laboratory studies often use different anesthetics when evaluating functional alterations in neurocircuitry. These anesthetics can influence glutamate neurotransmission through interactions with various molecular targets making it challenging to determine if observed differences are due to pathophysiology or anesthetic used. Thus, meta-analysis and literature reviews that do not consider anesthetic choice may result in inaccurate interpretations given the lack of information on the influence that anesthesia has on certain outcome measures of glutamate kinetics.

Despite the widespread use of various anesthetics, the fundamental question remains whether different anesthetics can influence neurochemical signaling and their mechanism of action. Urethane and isoflurane are commonly used anesthetics in translational neuroscience experiments, with ~8290 papers identified for urethane and isoflurane since 1981 (PubMed search for “urethane” AND “brain” or “isoflurane” AND “brain” from 1981 to 2020; Figure 1). In 1981, only five publications used isoflurane compared to 62 using urethane (7.5% isoflurane use). In 2020, 117 publications used isoflurane, and only 59 used urethane (66.5% isoflurane use). With this transition, it is important to consider whether specific outcome measures of experiments are influenced differentially between the two anesthetics so that researchers can build on previous bodies of knowledge. Furthermore, this knowledge would be a necessary consideration for experimental design and interpretation of future studies evaluating aspects of neuronal glutamatergic communication as we continue to apply them to understand neurochemical signaling under homeostatic and pathological conditions.

**Figure 1.**
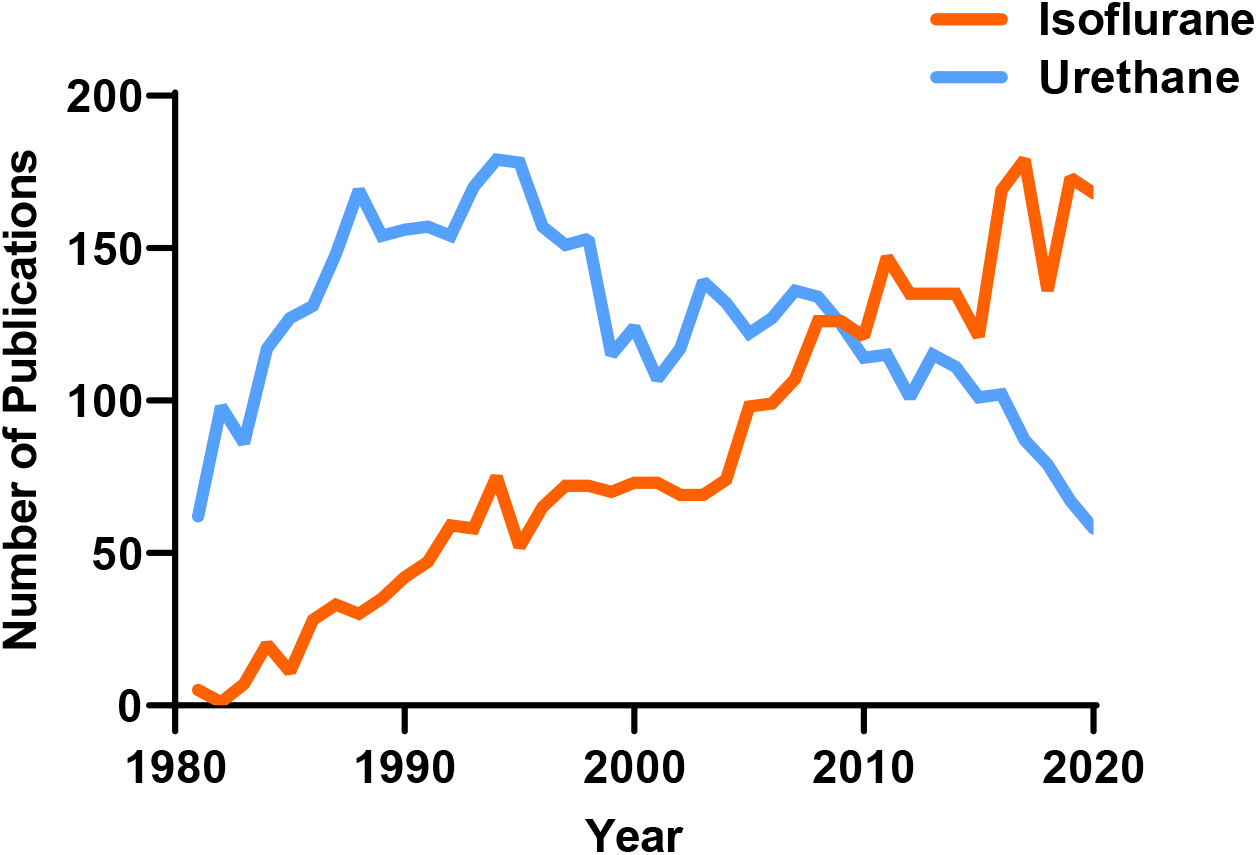
The number of publications using urethane and isoflurane for neuroscience research between 1981-2020. A PubMed search indicates the use of urethane has decreased while the use of isoflurane has increased over the past 40 years.

KCl-evoked glutamate overflow and glutamate clearance kinetics are commonly used metrics of glutamatergic communication, measured by microdialysis or amperometry in anesthetized rats. Both urethane and isoflurane are known to suppress glutamate neurotransmission but work through dissimilar mechanisms that could differentially influence glutamate signaling. Urethane, also known as ethyl carbamate, produces long-lasting, steady levels of anesthesia with reports indicating minimal effects on circulation, respiration, autonomic function, reflex responses, and GABA neurotransmission [2-6]. Urethane also induces alternating cycles of EEG activity that resemble REM sleep and wakefulness, indicating that brain activity is preserved to a certain extent [7]. Recent complementary studies indicate that urethane anesthetized rats retained functional connectivity patterns most similar to awake animals during resting-state functional magnetic resonance imaging in comparison to isoflurane [6]. While the mechanism of action is poorly understood, there is evidence that urethane primarily works through ion channels, distinct from isoflurane, influencing both inhibitory and excitatory systems [7]. These characteristics make urethane a popular anesthetic for electrophysiological experiments, but the actual influence on evoked glutamate release and clearance is unknown.

Given the benefits and ease of use with urethane, the reported carcinogenic properties of urethane primarily constrain urethane use to non-survival experiments [8, 9]. To use a more clinically relevant anesthetic, an increasing number of research papers using electrochemical and electrophysiological techniques have begun to use isoflurane. Isoflurane, a halogenated ether, has been implemented for its swift induction and recovery times and the precise control it allows over the amount of anesthesia delivered per unit time [10]. Mechanistically, evidence supports that isoflurane’s primary mechanism of action is mediated through the enhancement of GABAergic neurotransmission, thus inhibiting glutamate neurotransmission. In addition to different molecular mechanisms of action, urethane and isoflurane anesthesia have also been indicated to influence other brain regions to varying and opposing degrees. Paasonen et al. compared functional connectivity in several cortical and subcortical pathways under urethane and isoflurane protocols [5]. It was found that rats anesthetized with urethane retained similar connectivity in the cortical regions, but connectivity was suppressed in subcortical regions, including the hippocampus and thalamus. However, other studies comparing the influence of isoflurane and urethane on cortical networks in mice have shown the presence of stimulus-induced neuronal synchrony in both paradigms [11], highlighting a lack of congruency in previous reports where comparative studies are needed.

In this study, we sought to evaluate the relative comparability of KCl-evoked glutamate overflow and glutamate clearance kinetics collected via amperometric recordings from rats anesthetized by either isoflurane or urethane. Glutamate selective microelectrode arrays coupled with micropipettes filled with isotonic KCl solution or exogenous glutamate were placed within the cortex, hippocampus, or thalamus of naïve rats anesthetized with either isoflurane or urethane. The target regions were chosen because they are commonly implicated in cognition, somatosensation, and neuropathic pain.

## MATERIALS AND METHODS

### Subjects

Young adult male Sprague-Dawley rats (3-4 month old; 359-438 g upon delivery) were purchased (Envigo, Indianapolis, IN) and pair housed in disposable cages (Innovive, San Diego, CA) under normal 12:12 h light:dark cycle in a temperature- and humidity-controlled vivarium. Rats were provided food (Teklad 2918, Envigo, Indianapolis, IN) and water *ad libitum*. Group size estimates were determined from previous work [12], where *n* = 6-10/group could predict >90% power to detect a significant change in outcome measures. All procedures were conducted in accordance with the National Institutes of Health (NIH) Guidelines for the Care and Use of Laboratory Animals care and were approved by the University of Arizona College of Medicine-Phoenix Institutional Animal Care and Use Committee (protocol #18-384).

### Microelectrode Arrays

Ceramic-based MEAs encompassing four platinum recording surfaces (15 × 333 µm; S2 configuration) aligned in a dual, paired design were obtained from Quanteon LLC (Nicholasville, KY) for *in vivo* anesthetized recordings. MEAs were fabricated and selected for recordings using previously described measures of 0.125% glutaraldehyde and 1% GluOx [13-17]. MEAs were made glutamate selective as previously described [13, 18, 19]. Prior to *in vivo* recordings, a size exclusion layer of m-phenylenediamine dihydrochloride (Acros Organics, NJ) was electroplated to all four platinum recording sites with the use of the FAST16 mkIII system (Fast Analytical Sensor Technology Mark III, Quanteon, LLC, Nicholasville, KY) to block potential interfering analytes such as ascorbic acid, catecholamines and other indoleamines [20]. The four platinum recording sites consisted of two glutamate-sensitive and two sentinel channels. The glutamate sensitive-channels were coated with 1% BSA, 0.125% glutaraldehyde and 1% glutamate oxidase.

### Microelectrode Array Calibration

MEAs were calibrated with a FAST16 mkIII system to accurately record glutamate concentrations through the creation of a standard curve for each coated MEA within 24 hours of use in *in vivo* recordings, as described previously [18]. Briefly, the MEAs were submerged into 40 ml of stirred 0.05 M phosphate-buffered saline (PBS) solution kept in a water bath and allowed to reach a stable baseline before beginning the calibration.

Aliquots from stock solutions were added in succession following equilibrium such that 500 µL of ascorbic acid, three additions of 40 µL of L-glutamate, two additions of 20 µL of dopamine, and 40 µL of H_2_O_2_ were added to produce final concentrations of 250 µM of ascorbic acid, 20, 40 and 60 µM of L-glutamate, 20 µM and 40 µM dopamine and 8.8 µM of H2O2 respectively in the beaker of PBS (pH 7.4). A representative calibration is depicted in Figure 2. From the MEA calibration, the following metrics were calculated and used for the determination of inclusion within the *in vivo* recordings: slope (sensitivity to glutamate), the limit of detection (LOD) (lowest amount of glutamate to be reliably recorded), and selectivity (ratio of glutamate to ascorbic acid). For the study, a total of 48 MEAs were used with 96 recording sites (Slopes: >5 pA/µM, LOD: <2.5 µM, and selectivity: >20:1). A representative MEA calibration is shown in Figure 2.

**Figure 2.**
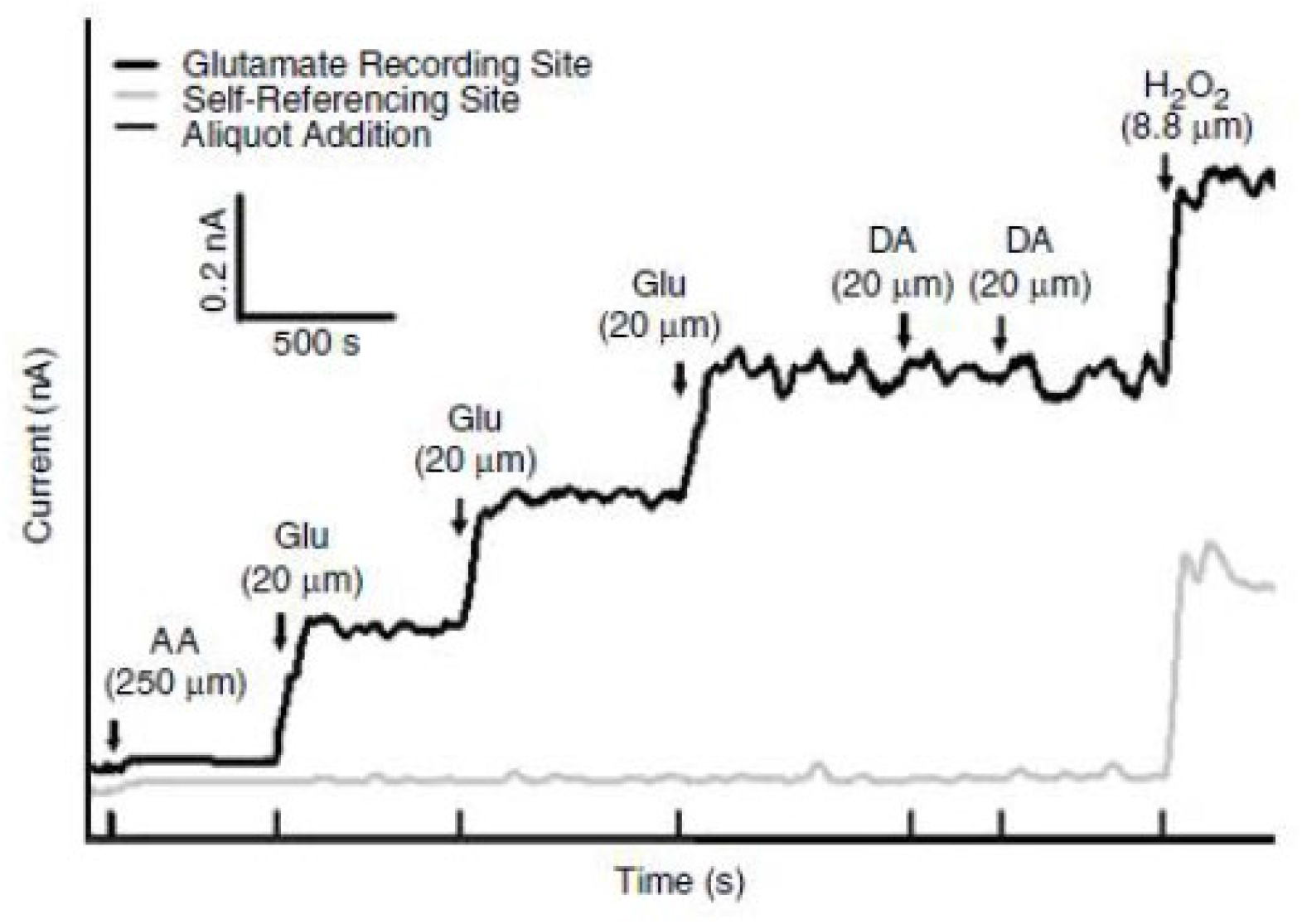
Representative calibration of a glutamate selective microelectrode array (MEA). Representative MEA calibration prior to *in vivo* recordings in which only one glutamate selective recording site and one self-referencing site are represented. Aliquots of 250 µM ascorbic acid (AA), 20 µM glutamate (Glu), 2 µM dopamine (DA), and 8.8 µM H2O2 are represented by vertical bars on the x-axis.

### Microelectrode Array/Micropipette Assembly

Following calibration, a single-barrel glass micropipette was attached to the MEA with the following steps to allow for the local application of solutions during *in vivo* experiments. A single-barreled glass capillary with a filament (1.0 × 0.58 mm^2^, 6” A-M Systems, Inc., WA) was pulled to a tip using a Kopf Pipette Puller (David Kopf Instruments, CA). The pulled pipette was then bumped using a microscope and a glass rod to have an inner diameter averaging 11.1 *µ*m ± 0.56. The pulled glass pipettes were embedded in modeling clay and attached to the circuit board above the MEA tip. Molten wax was applied to the embedded pipette to secure the MEA/micropipette assembly and prevent its movement during the recording. The pipette attachment was performed under a microscope to carefully place the tip of the pipette above the glutamate sensitive sites from the surface of the electrode. Measurements of pipette placement were confirmed using a microscope with a calibrated reticule in which the pipette was approximately 45 to 105 µm from the electrode sites (averaging 78.7 µm ± 4.9).

### Reference Electrode Assembly

Silver/silver chloride reference electrodes were fabricated from Teflon-coated silver wire to provide an *in vivo* reference for the MEA. A 0.110-inch Teflon-coated silver wire (A-M Systems, Carlsborg, WA) was prepared by stripping approximately ¼” of Teflon coating from each end of a 6” section, soldering one end to a gold-plated socket (Ginder Scientific, Ottawa, ON) and the other being prepared to be coated with silver chloride. This end was placed into a 1 M HCl saturated with NaCl plating solution. A 9V current was applied to the silver wire (cathode) versus the platinum wire (anode) for approximately 5 minutes. Upon completion, the silver/silver chloride reference electrode was placed in a light-sensitive box until implanted.

### Surgery

Rats were randomly assigned to receive either isoflurane or urethane until the pedal reflex was eliminated. Isoflurane was delivered via a nose cone (5% induction and 2% maintenance in oxygen) throughout the recordings. 25% urethane (Sigma Aldrich, St. Louis, MO) was administered as serial intraperitoneal injections (1.25-1.5 g/kg). Following the loss of the pedal response, each rat was then placed into a stereotaxic apparatus (David Kopf Instruments) with nonterminal ear bars. Body temperature was maintained at 37 °C with isothermal heating pads (Braintree Scientific, MA). A midline incision was made in which the skin, fascia, and temporal muscles were reflected, exposing the scalp. A bilateral craniectomy was then performed using a Dremel, exposing the stereotaxic coordinates for the somatosensory cortex, hippocampus, or thalamus. Dura was removed prior to the implantation of the MEA. Brain tissue was hydrated by applying saline-soaked cotton balls and gauze. Using blunt dissection, a silver/silver chloride-coated reference electrode wire was placed in a subcutaneous pocket on the dorsal side of the rat [21, 22]. All experiments were performed during the light phase of the 12 h-dark/light cycle.

### *In vivo* Amperometric Recordings

Amperometric recordings performed here were similar to previously published methods [13, 23, 24]. Briefly, a constant voltage was applied to the MEA using the FAST16 mkIII recording system. *In vivo* recordings were performed at an applied potential of +0.7 V compared to the silver/silver chloride reference electrode wire. All data were recorded at a frequency of 10 Hz and amplified by the headstage piece (2 pA/mV). Immediately prior to implantation of the glutamate-selective MEA-pipette assembly, the pipette was filled with 120 mM of KCl (120 mM KCl, 29 mM NaCl, 2.5 mM CaCl_2_, pH 7.2 to 7.5) or 100 µM L-glutamate (100 µM L-glutamate in 0.9% sterile saline pH 7.2-7.5) immediately prior to implantation and *in vivo* recording. The concentrations for both solutions have been previously shown to elicit reproducible potassium-evoked glutamate overflow or exogenous glutamate peaks [15, 25, 26]. Solutions were filtered through a 0.20 µm sterile syringe filter (Sarstedt AG & Co. Numbrecht, Germany) attached to a 1 mL syringe with a 4-inch, 30-gauge stainless steel needle with a beveled tip (Popper and Son, Inc, NY) while filling the micropipette. The open end of the micropipette end was then connected to a Picospritzer III (Parker-Hannin Corp., General Valve Corporation, OH) with settings to dispense fluid through the use of nitrogen gas in nanoliter quantities as measured by a dissecting microscope (Meiji Techno, San Jose, CA) with a calibrated reticule in the eyepiece [27, 28].

Once the MEA-micropipette apparatus was securely attached to the Picospritzer and FAST system, bregma was measured using an ultraprecise stereotaxic arm. MEA-micropipette constructs were implanted in the cortex (AP, -2.8 mm; ML, ±5.0 mm; DV, -1.0 mm vs. bregma), hippocampus (AP, -3.5 mm; ML, ±3.0 mm; DV, -2.6 to -3.75 mm vs. bregma) and thalamus (AP, -3.5 mm; ML, ±3.0 mm; DV, -5.6 mm vs. bregma) based on the coordinates from Paxinos and Watson [29] (Figure 3A). Glutamate and KCl-evoked measures were recorded in both hemispheres in a randomized and balanced experimental design to mitigate possible hemispheric variations or effects of anesthesia duration.

**Figure 3.**
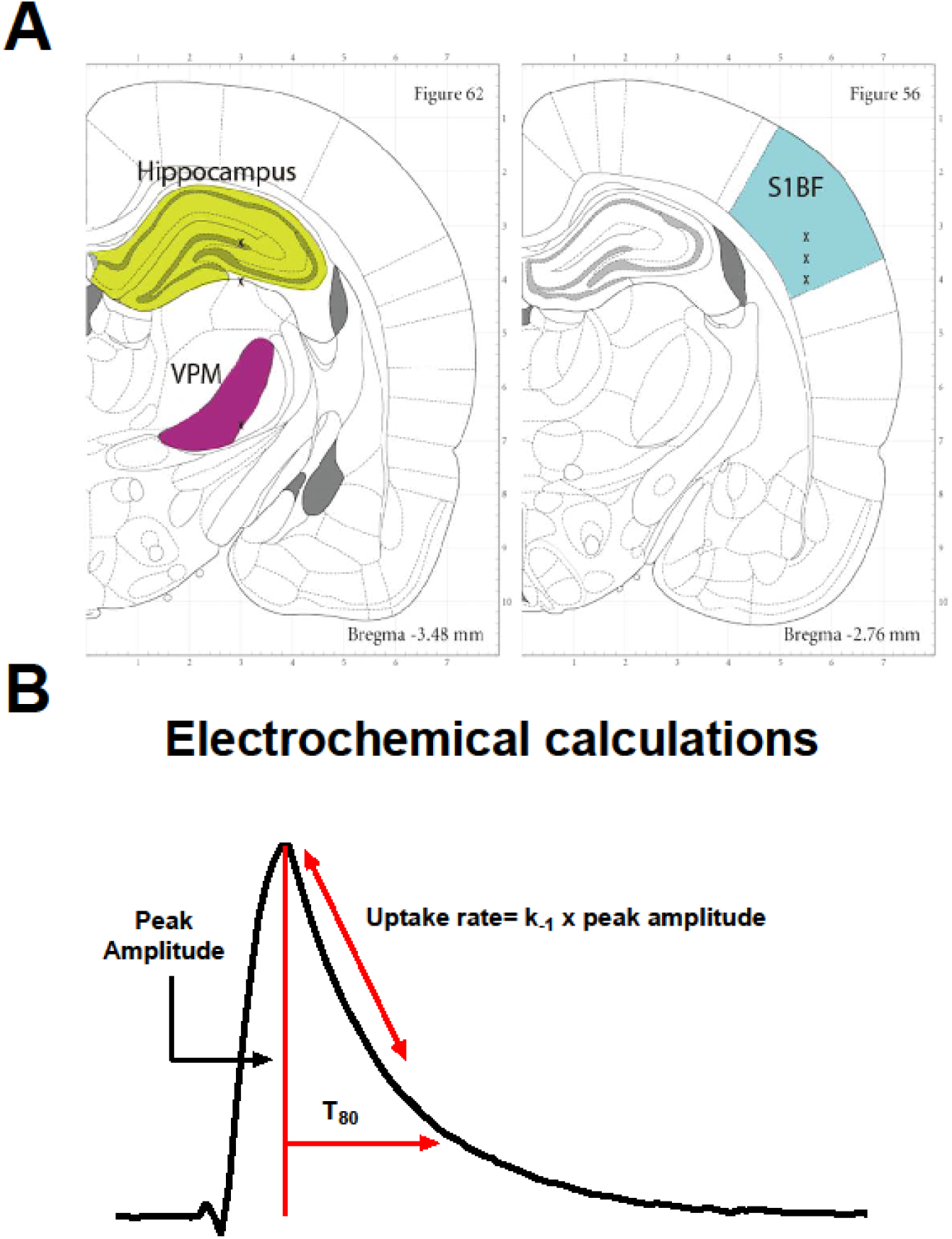
Recording regions and amperometric calculations. **(A)** Anatomical regions of interest (ROI) in the rodent brain were the hippocampus (green), thalamus (purple), and cortex (blue). Image modified from Paxinos and Watson (2007). (**B)** Representative peak, showing glutamate concentration (µM) as a function of time in seconds. Amperometric calculations used in the analysis (peak amplitude, uptake rate, and T_80_) are shown.

### KCl-Evoked Overflow of Glutamate Analysis Parameters

All recordings were conducted at 10 Hz and analyzed without signal processing or filtering the data. Once the glutamate MEA was lowered, and the electrochemical signal had reached stable baseline, pressure injections of a known volume of 120 mM potassium was locally applied to induce depolarization and subsequent overflow of glutamate from synapse was measured by the MEA. The volume of each application of KCl was predetermined to ensure maximal response (largest amount of glutamate released) which was confirmed by a smaller evoked peak following the 2-minute interval. The maximal amplitude of the glutamate response (µM) of the first peak was the primary outcome measure for analysis.

### Glutamate Clearance Analysis Parameters

All recordings were conducted at 10 Hz and analyzed without signal processing or filtering of the data. Once the electrochemical signal had reached a baseline, 100 µM glutamate was locally applied into the extracellular space. Additions of exogenous glutamate were applied at 30-second intervals for reproducible glutamate peaks. Clearance parameters of 3 amplitude-matched peaks were averaged to create a single representative value per recorded region per rat. Primary outcome measures for analysis include the uptake rate and the time taken for 80% of the exogenous glutamate to clear the extracellular space (T_80_). The uptake rate is calculated by multiplying the uptake rate constant (k_-1_) by the peak’s maximum amplitude. Diagrammatic example of these calculations is shown in Figure 3B.

### MEA Placement Verification

Immediately following *in vivo* anesthetized recordings, rats were transcardially perfused with PBS and post-fixed in 4% paraformaldehyde (PFA). Brains were cryoprotected, sectioned, and stained with a hematoxylin and eosin stain to confirm MEA electrode placement. No electrode tracts were excluded due to placement (data not shown).

### Statistical Analysis

The amperometric data were saved on the FAST 16 mkIII system. Datasets were analyzed with FAST Analysis software (Jason Burmeister Consulting) and processed using a customized Microsoft Excel^®^ spreadsheet.

Inclusion criterion for data analysis of KCl-evoked response was maximum amount of glutamate overflow. Inclusion criterion for glutamate clearance kinetics was in the amplitude of 10 to 23 µM to control for Michaelis-Menten kinetics. All data are presented in the form of mean + standard error mean (mean ± SEM) and analyzed using the statistical software GraphPad Prism 9.4.1. Comparisons between anesthetics used a two-tailed Student’s *t*-test. Differences between regions were determined using a one-way ANOVA with Tukey’s post-hoc comparison. All data sets were evaluated for variance (Brown-Forsythe) and normality (Kolmogorov-Smirnov) to ensure assumptions are met for analysis. Outliers were evaluated using a ROUT method with a false discovery rate of (Q) 1%. Differences were considered statistically significant when p < 0.05.

## RESULTS

### Levels of evoked release of glutamate were similar in urethane and isoflurane-anesthetized rats

Surrounding neuronal tissue was depolarized with volume-matched (75-150 nL) applications of 120 mM isotonic KCl, and the maximum amplitude of glutamate was recorded. Cortical recordings revealed no significant differences in the amount of glutamate overflow between rats anesthetized with isoflurane or urethane (*N* = 6-7 rats; 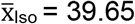 µM, 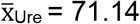 µM; *t*_11_ = 1.629; p = 0.131 µM; Figure 4A). Similarly, KCl-evoked glutamate overflow was independent of anesthetics in the hippocampus (*N* = 7 rats; 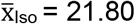 µM, 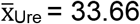 µM; *t*_12_ = 1.247; p = 0.236; Figure 4B) or thalamus (*N* = 8 rats; 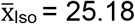 µM, 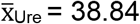 µM; *t*_14_= 1.299; p = 0.214; Figure 4C).

**Figure 4.**
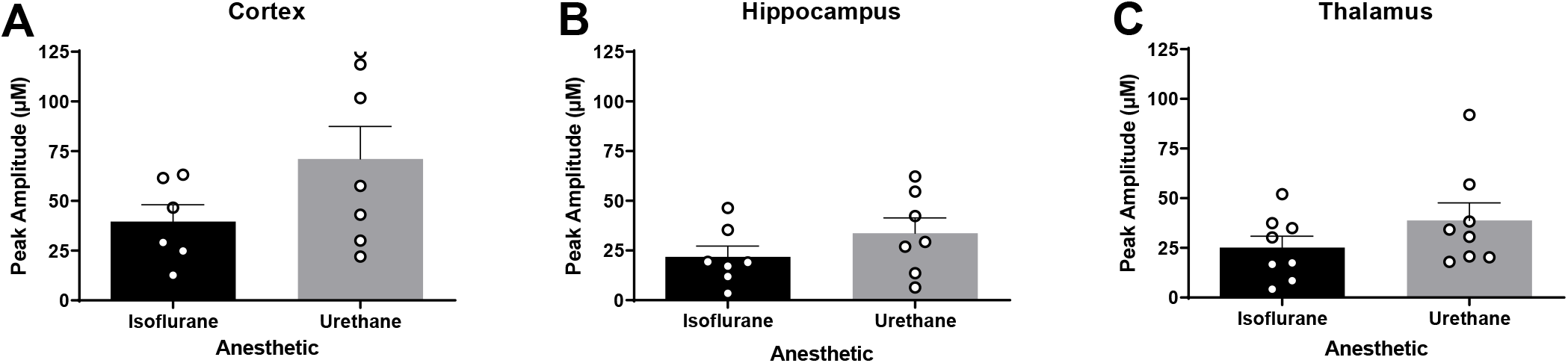
Levels of evoked release of glutamate were similar in urethane and isoflurane-anesthetized rats. Local applications of volume-matched 120 mM potassium chloride (KCl) were made to the cortex, hippocampus, and thalamus. There was no significant difference between the glutamate overflow measured for rats anesthetized with isoflurane or urethane in the **(A)** cortex (*t*11 = 1.629, p = 0.131), (**B)** hippocampus (*t*_12_ = 1.247, p = 0.236), or (**C)** thalamus (*t*_14_ = 1.299, p=0.214). Bar graphs represent mean + SEM.

### Glutamate uptake rate was faster in urethane anesthetized rats

In each region of interest (ROI), glutamate clearance from the extracellular space was evaluated. Local applications of 100 µM exogenous glutamate were amplitude-matched at the time of administration. Cortical recordings revealed no significant differences in uptake rate between rats anesthetized with isoflurane or urethane (*N* = 6-7 rats; 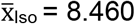 µM/sec., 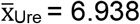 µM/sec.; *t*_11_ = 0.544; p = 0.597; Figure 5A). Uptake rate was also similar between anesthetics in the hippocampus (*N* = 7 rats; 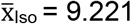 µM/sec., 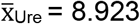 µM/sec.; *t*_12_ = 0.148; p = 0.884; Figure 5B). However, the thalamus of urethane anesthetized rats showed significantly faster uptake rate in comparison to isoflurane-anesthetized rats (*N* = 6-9 rats; 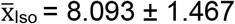 ± 1.467 µM/sec., 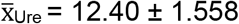 = 12.40 ± 1.558 µM/sec.; *t*_13_ = 2.817; p = 0.0145; Figure 5C). No differences were identified in the cortex (*N* = 5-7 rats; 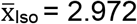 seconds, 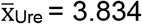 seconds; *t*_10_ = 1.329; p = 0.213; Figure 6A), hippocampus (*N* = 5-7 rats; 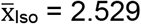 seconds, 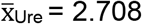 seconds; *t*_9_ = 0.525; p = 0.612; Figure 6B), or thalamus (*N* = 6-9 rats; 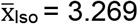 seconds, 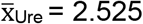 seconds; *t*_13_ = 1.827; p = 0.090; Figure 6C).

**Figure 5.**
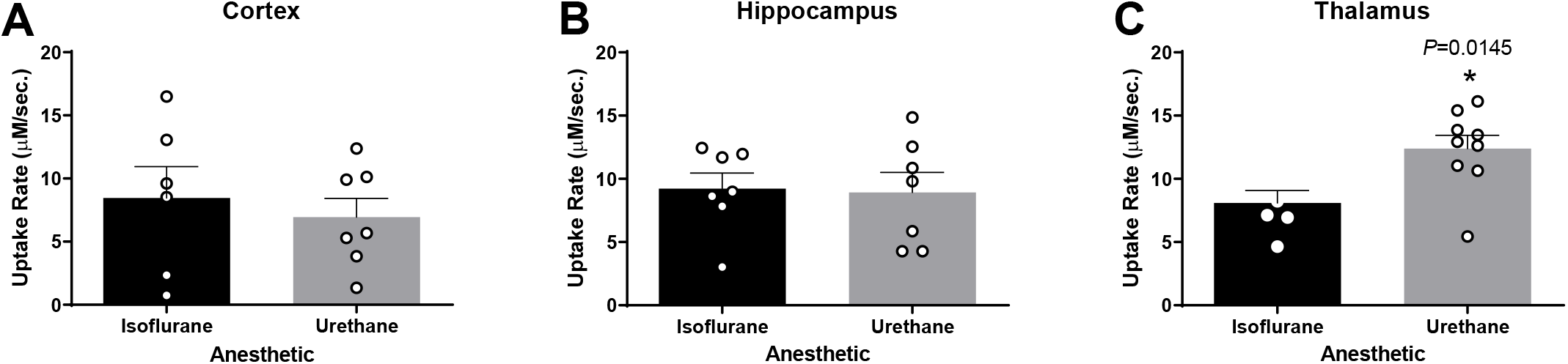
Glutamate uptake rates were similar in the cortex and hippocampus and different in the thalamus of urethane and isoflurane-anesthetized rats. Amplitude-matched signals from local applications of 100 µM exogenous glutamate compared for extracellular glutamate clearance in the cortex, hippocampus, and thalamus. No significant difference in the uptake rate between rats anesthetized with isoflurane or urethane in the (**A)** cortex (*t*_11_ = 0.544, p = 0.597) and **(B)** hippocampus (*t*_12_ = 0.148, p = 0.884). **(C)** Urethane administration was associated with a significantly faster uptake rate in thalamus than under isoflurane exposure (*t*_13_ = 2.817, p = 0.0145). Bar graphs represent mean + SEM.

**Figure 6.**
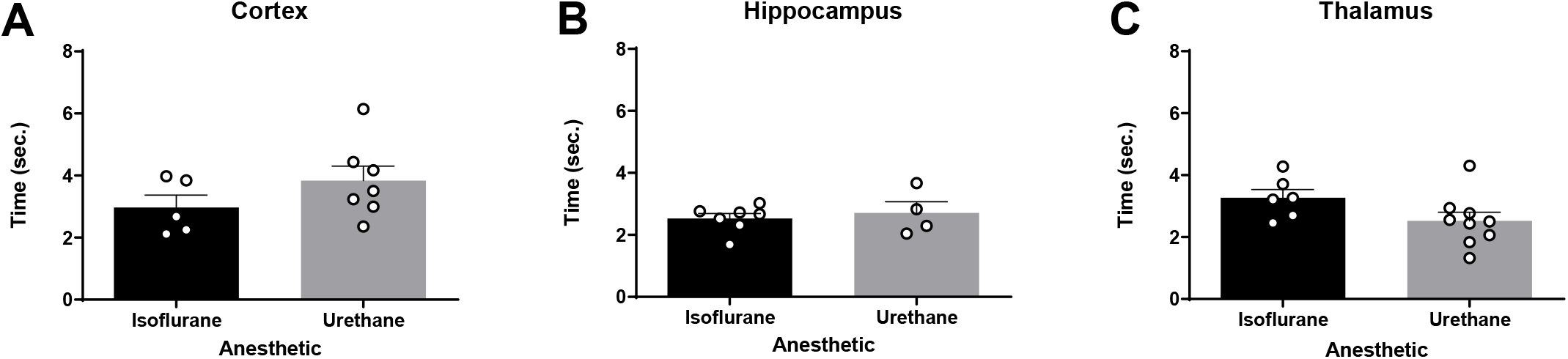
Extracellular clearance times (T_80_) were similar in urethane and isoflurane-anesthetized rats. Local application of 100 µM exogenous glutamate resulted in amplitude-matched signals to assess the T_80_ in the cortex, hippocampus, and thalamus. There was no significant difference in the time taken for 80% of the maximal amplitude to clear between rats anesthetized with isoflurane or urethane in the **(A)** cortex (*t*_10_ = 1.329, p = 0.213), **(B)** hippocampus (*t*_9_ = 0.525, p = 0.612), or **(C)** thalamus (*t*_13_ = 1.827, p = 0.090). Bar graphs represent mean + SEM.

### Urethane anesthesia influences brain region dependent differences of glutamate kinetics

Aspects of glutamate neurotransmission were compared between brain regions under the influence of single anesthetic to determine region specific differences. No significant differences were detected between the cortex, hippocampus, and thalamus when using isoflurane for KCl-evoked glutamate overflow, uptake rate, and T_80_ (Figure 7A-C). Additionally, no differences were detected between regions for KCl-evoked overflow (Figure 7D) when using urethane anesthesia. Region dependent differences were detected in glutamate clearance with both the uptake rate (F_2,20_ = 4.41; p = 0.026; Figure 7E) and T_80_ (F_2,14_ = 3.79; p = 0.044; Figure 7F) being statistically significant between the thalamus and cortex recordings. Uptake rate in the thalamus was significantly faster than the cortex (p = 0.023), with no differences detected between the cortex or thalamus and the hippocampus (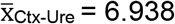 µM/sec, 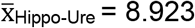 µM/sec, 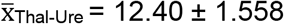 ± 1.558 µM/sec). Faster uptake rate in the thalamus was supported by a shorter time to clear 80% of the glutamate signal in comparison to the cortex (p = 0.042). There were no differences detected between the T_80_ in the cortex or thalamus compared to the hippocampus 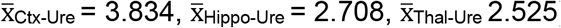.

**Figure 7.**
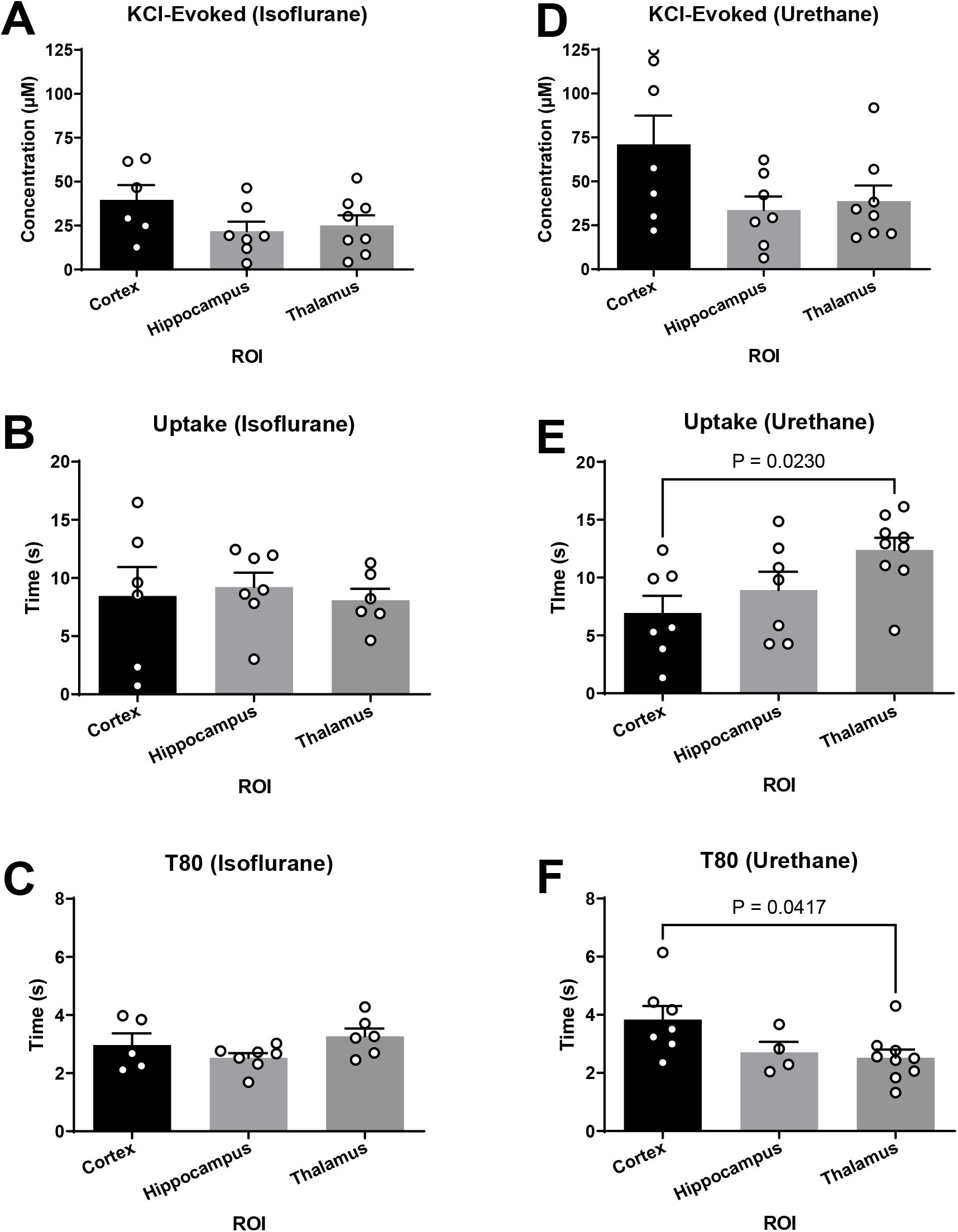
Glutamate clearance under urethane was capable of distinguishing region-dependent differences. **(A-C)** Under isoflurane, all outcome measures were similar between the cortex, hippocampus, and thalamus. **(D)** No significant differences were detected in KCl-evoked glutamate release across regions when using urethane. **(E-F)** Glutamate clearance kinetics under urethane changes as a function of the brain region when evaluating uptake rate (F_2,20_ = 4.41; p = 0.02) and T_80_ (F_2,14_ = 3.79; p = 0.04), where uptake rate was significantly faster and clearance time was significantly shorter in the thalamus compared to the cortex. Bar graphs represent mean + SEM.

## DISCUSSION

These experiments were designed to test whether KCl-evoked glutamate overflow and glutamate clearance kinetics in the cortex, hippocampus, and thalamus were similar in rats anesthetized with either urethane or isoflurane. No differences in KCl-evoked glutamate overflow were identified by evaluating the maximal amplitude of glutamate released. The uptake rate and clearance time (T_80_) of locally applied exogenous glutamate were also similar under both anesthetics in the cortex and hippocampus, however, the uptake rate under urethane was significantly faster in the thalamus compared to isoflurane. Further, urethane treatment allowed the capability of measuring region-specific differences, where all regions were similar with isoflurane anesthetization. These data support the need for careful interpretation when comparing recordings from rodents under urethane or isoflurane anesthesia as certain parameters of glutamate uptake vary based on choice of anesthesia and brain region.

Local application of KCl causes a net positive of the resting membrane potential of surrounding neurons and glia, which results in subsequent action potentials that release stored neurotransmitters. The glutamate selective MEAs record sub-second changes in glutamate overflow into the extracellular space, such that the maximum amplitude can be statistically evaluated. Measured values are indicative of the amount of glutamate possible to be released during synaptic signaling. Alterations of values obtained from KCl-evoked glutamate overflow may result from changes to presynaptic neuron release, glutamate output from surrounding glia, decreased glutamate clearance, and the changes to glutamate transporters (GLT-1, GLAST, EAAC), GABAergic or modulatory neurotransmission, or alterations to mGluR regulation of glutamate release. [13, 23, 30-32].

Local application of exogenous glutamate enables the study of glutamate clearance kinetics in the extracellular space. The main contributors to the clearance of glutamate from the extracellular space in the cortex, hippocampus, and thalamus are astrocytic glutamate transporters EAAT1 (GLT-1; cerebral cortex: 90%; thalamus: 54%; data in comparison to the hippocampus) and EAAT2 (GLAST; cerebral cortex: 33%; hippocampus: 35%; thalamus: 22%; data in contrast to the cerebellum), and to a lesser extent, post-synaptic transporter EAAT3 [1, 33, 34]. Changes in glutamate clearance may indicate the function of glutamate transporters, transporter trafficking, transporter capacity, and transporter affinity [35]. Extracellular clearance parameters, including uptake rate, represent the velocity of the transporters, while T_80_ indicates the affinity of glutamate to the transporters [36]. Previous work has shown that alterations to the location (trafficking) or expression of GLT-1 and GLAST correlate to changes in glutamate clearance parameters as well [37, 38]. Conflicting literature evaluating the influence of isoflurane on glutamate uptake in *in vitro* models indicates enhancements in a dose-dependent manner [39-41], suppression/inhibition [42, 43], or no change [44]. Further, isoflurane use has been shown to increase the surface-level expression of EAAT3 [45]. While studies addressing the influence of urethane on glutamate transporters are lacking, the present data indicate that urethane interacts with the neurotransmitter system in the thalamus differently, such that regional dependent differences in glutamate clearance can be detected. Perhaps cellular, laminar, and regional heterogeneities of glutamate transporter distribution in the thalamus are contributing to this effect [46]. However, additional studies are needed to confirm whether this effect is due to the anesthetics having a differential influence on transporter function/expression or whether other distinct mechanisms of action culminate in minor region-dependent clearance kinetics.

The pharmacological mechanisms for anesthesia by urethane and isoflurane need to be better understood at the neurochemical level, making it difficult to predict the effect on aspects of neurotransmission [47]. Both urethane and isoflurane have been indicated to dampen overall glutamate neurotransmission, reported by evidence of less glutamate released, altered reuptake, reduced cellular excitability, decreased evoked glutamate overflow, and decreased tonic levels [2, 42, 48-53]. These effects may be mediated through the interaction between urethane and isoflurane with various GABAergic and glutamatergic receptors, as well as similar antagonistic effects at NMDA and AMPA receptors [54, 55]. Local field potential values were also similar under both anesthetics [47]. While each respective anesthetic’s molecular mechanism remains only partially understood, similarities amongst molecular targets and aspects of neuronal communication may contribute to comparable alterations in glutamate neurotransmission. Alternate sites of action of the anesthetics indicate a dose-dependent influence on factors that mediate glutamate neurotransmission [54, 56]. Isoflurane has been shown to bind to GABAa receptors [57], glutamate receptors, and glycine receptors, inhibit potassium channels [58, 59], and hyperpolarize neurons [60]. Hara *et al*. describes several unique molecular targets of urethane that influence glutamate neurotransmission, including enhancement of GABAa and glycine receptors, while various NMDA and AMPA subtypes are inhibited dose-dependently [54]. Furthermore, a 30% decrease in glutamate concentrations in the cerebral cortex of rats under urethane exposure is observed compared to un-anaesthetized rats [2]. It is indicated that urethane and isoflurane suppress presynaptic glutamate release associated with increased excitatory postsynaptic currents (EPSCs) in the hippocampus and thalamus, respectively [3, 61]. Further, it is shown that glutamate neurotransmission is reduced in cortical inhibitory interneurons with urethane exposure [62]. These reports indicate that other parameters, like basal extracellular levels, electrophysiological characteristics, and cell types not evaluated in these experiments, could be disrupted. Importantly, a recent study indicated that both isoflurane and urethane exposure had no significant impact on the neuronal response to whisker stimulation in mice [13, 18, 63, 64].

Valuable data may be obtained using neurochemical analysis under anesthesia. Whether accomplished through amperometry, microdialysis, or electrophysiology, a better understanding of neuronal communication helps identify pathological changes in glutamate neurotransmission within behaviorally relevant circuits [65]. Additionally, animal-specific factors (species, strain, age, and sex) influence glutamate neurotransmission, and the precise mechanism (or combination of mechanisms) that culminates in overall dampened glutamate neurotransmission remains theoretical and may confound biological relevance [13, 18, 37, 66-71]. Accordingly, our data indicate the use of either isoflurane or urethane results in similar, comparable effects on glutamatergic parameters in the cortex and hippocampus, despite being accomplished through dissimilar mechanisms. However, other functional outcomes of neuronal communication under these anesthetics have not been shown to result in a similar effect. Indeed, it is reported that anesthetic doses of isoflurane dampen sub-cortical activity inducing synchronous cortico-striatal fluctuations, while the use of urethane resulted in recordings most similar to awake rodents [5]. Neuronal activity measured utilizing BOLD fMRI reported that isoflurane and urethane dampens responses when compared to unanesthetized subjects but does not comment on which anesthetic is more comparable to the awake state [72-74]. Studies investigating changes in molecular components that are known to interact with a given anesthetic must use caution when interpreting results and attributing changes to a specific mechanism. These and other studies indicate that not all neuronal circuits respond identically under anesthesia. Thus, investigating anesthesia’s effect on other neuromodulators (5-HT, GABA, DA, ACh) and their functional outputs in specific circuits should be considered when interpreting comparisons across several studies. It is even recommended that studies evaluating physiologic parameters controlled by the CNS should use caution when measuring their output while under anesthesia since CNS-mediated physiologic functions may not result in comparable changes under isoflurane or urethane despite PCO2, O2, pH, and heart rate having been shown to be similarly affected [5]. Therefore, experiments should be carried out in preparation for moving into awake-behaving models, where simultaneous recordings of behavior and glutamate signaling are possible. In doing so, we can discover aberrant glutamate signaling and determine the efficacy of therapeutic approaches relevant to treat several brain disorders marked by maladaptive behaviors.

The safety and efficacy of anesthetic on animals has historically influenced researchers’ choice when considering study design. Urethane is classified by the International Agency for Research on Cancer as a group 2A carcinogen and thus considered “probably carcinogenic to humans.” This designation has limited urethane use primarily to non-survival studies despite its ease of use and favorable profile. In doing so, using urethane as a sole anesthetic in longitudinal studies with multiple data collection time points in the same animals is not considered humane for the risk of tumor formation. However, the inclusion of urethane as part of an anesthetic cocktail provides researchers with significant advantages. In mice, it was found that using 560 mg/kg alongside ketamine and xylazine was not carcinogenic [75]. Ultimately this cocktail was advantageous as the use of low doses of all three anesthetics limited confounding changes in electrical potentials observed when a ketamine/xylazine combination is used alone [75-77]. Of note, these studies provide evidence that the combination of multiple anesthetics leads to the capitalization of favorable effects.

## Conclusions

In conclusion, these data support the hypothesis that isoflurane and urethane similarly influence KCl-evoked glutamate overflow and glutamate clearance parameters. Electrochemical recordings from outcome measures under urethane or isoflurane anesthesia with a similar protocol, inclusion, and exclusion criterion can be integrated into interpretations with careful regard to other potential influencing factors, especially region of interest. Finally, future studies evaluating glutamate neurotransmission should focus on the beneficial aspects of anesthetic when optimizing the experimental design as well as the inclusion of the female sex to evaluate any sex-specific sensitivity.

## List of abbreviations

*AD*: Alzheimer’s disease
*ADHD*: attention-deficit/hyperactivity disorder
*CNS*: central nervous system
*HD*: Huntington’s disease
*KCl*: potassium chloride
*LOD*: limit of detection
*MEAs*: microelectrode arrays
*PBS*: phosphate buffered saline
*PD*: Parkinson’s disease
*PFA*: paraformaldehyde
*ROI*: region of interest

## Declarations

## Ethics approval and consent to participate

Not applicable

## Consent for publication

Not applicable

## Availability of data and materials

All data that support the findings of this study are available from the corresponding author upon request.

## Authors’ contributions

J.A.B., G.K., and T.C.T. drafted and revised the manuscript, figures, and legends. They all independently analyzed the data and contributed to the interpretation of the data. J.A.B., G.K., and C.E.B. carried out the experiments. T.C.T. designed the research and acquired funding The content is solely the responsibility of the authors and does not necessarily represent the official views of the National Institutes of Health.

## Acknowledgments

We thank Carol Haussler for critical review of the manuscript.

